# Inhibition of the mitochondrial pyruvate carrier attenuates the integrated stress response activation in a cellular model of Huntington’s disease

**DOI:** 10.64898/2026.01.22.701013

**Authors:** Ângela Oliveira, Liliana M. Almeida, Jorge M. A. Oliveira, Brígida R. Pinho

## Abstract

Mitochondrial pyruvate carrier (MPC) inhibition was found protective in models of neurodegenerative diseases, such as Alzheimer’s and Parkinson’s. However, little is known about MPC as a potential therapeutic target in Huntington’s disease (HD), a neurodegenerative disorder with dysregulation of the pro-survival pathway integrated stress response (ISR). Here, we investigate if MPC inhibition modulates the ISR and mitigates mutant huntingtin (mut-Htt) proteotoxicity in a cellular HD model. We treated cells expressing N-terminal fragments of wild-type- (wt-) or mut-Htt with two MPC inhibitors (mitoglitazone and UK5099) or solvent control. Metabolism was assessed analysing resazurin reduction, oxygen consumption, extracellular acidification, and ATP levels. ISR activation and huntingtin proteostasis were assessed using western-blot and filter-trap assays. Mut-Htt-expressing cells showed decreased resazurin reduction and ATP levels, and increased eIF2*α* phosphorylation, indicating metabolic stress and ISR activation. MPC inhibitors (100 µM) increased resazurin reduction and decreased respiration. The latter was rescued by the membrane-permeant methyl pyruvate, which bypasses MPC inhibition. In wt-Htt-expressing cells, MPC inhibitors increased levels of ATP and ISR markers, suggesting metabolic adaptation and ISR activation. In mut-Htt-expressing cells, MPC inhibitors preserved ATP levels and attenuated mut-Htt-induced eIF2*α* phosphorylation but without changing soluble or aggregated mut-Htt levels. This work showed that MPC inhibition differentially modulates the ISR: it activates ISR in control cells and attenuates overactive ISR in mut-Htt-expressing cells. However, MPC inhibition did not impact the proteostasis of N-terminal fragment mut-Htt. Further studies are essential to explore MPC inhibition in less severe full-length mut-Htt-expressing models to better understand its therapeutic potential in HD.

## Introduction

Neurodegenerative diseases are a major societal challenge, aggravated by the current lack of disease-modifying treatments [1]. Previous studies have identified the mitochondrial pyruvate carrier (MPC) as a potential therapeutic target in neurodegenerative diseases [2-4].

MPC locates at the inner mitochondrial membrane and mediates pyruvate transport into mitochondria. Pharmacological inhibition of the MPC promoted dopaminergic neuron survival and improved locomotion in animal models of Parkinson’s disease (PD) [3, 4]. In Alzheimer’s disease (AD), pharmacological MPC inhibition for 3 months preserved glucose metabolic rate in the cerebellum of patients, whereas placebo-treated patients showed a significant decline [5]. Also, in an AD mouse model, astrocyte-specific deletion of the MPC1 subunit reduced amyloid and tau accumulation [2]. Mechanistically, reduction of MPC activity triggers metabolic reprogramming to compensate for the decreased pyruvate import into mitochondria, which is essential for the maintenance of the tricarboxylic acid (TCA) cycle. Indeed, MPC inhibition favours the consumption of glutamine, branched-chain amino acids and fatty acids [6, 7]. However, the mechanisms linking these mitochondrial metabolic changes to the observed neuroprotective effects remain unclear [8].

Metabolic changes, such as depletion of cytosolic amino acid pool, can activate the integrated stress response (ISR), a pro-survival homeostatic pathway, via the ISR kinase GCN2. GCN2 phosphorylates the eukaryotic initiation factor 2 alpha (eIF2α), a key effector of the ISR pathway [9, 10]. Phosphorylated eIF2α reduces global protein synthesis while promoting the selective expression of transcription factors such as activating transcription factor 4 (ATF4) or C/EBP-homologous protein (CHOP), which regulate the expression of proteins involved in amino acid transport and synthesis, autophagy and cell death pathways [9]. Previous research has demonstrated that MPC inhibition triggers ISR activation in human hair follicles [11] and in cellular models of prostate [12] and lung [13] cancer, while attenuating ISR activation induced by endoplasmic reticulum (ER) stressors thapsigargin or tunicamycin in primary hepatocytes and myoblasts [14]. These results highlight the need for further research to clarify the role of MPC inhibition in the ISR pathway and if ISR modulation may contribute to the neuroprotective effects of MPC inhibition.

HD is a monogenic neurodegenerative disease caused by an expanded polyglutamine (polyQ) stretch in the huntingtin (Htt) protein [15, 16]. This polyQ expansion increases the propensity of mutant Htt (mut-Htt) for proteolysis, misfolding and aggregation [17, 18]. Mut-Htt-mediated toxicity includes impairment of energy metabolism [19], ER stress [20], collapse of the protein homeostasis (proteostasis) network [18] and dysregulation of the ISR [21]. According to a recent review, the involvement of the MPC in HD pathophysiology has not yet been investigated [8]. Moreover, as far as we could find in the literature (Scopus search: (MPC OR “Mitochondrial pyruvate carrier”) AND huntingt*), the therapeutic potential of MPC inhibition in HD has not been investigated. Thus, it remains unknown whether MPC inhibition can alter mut-Htt levels and aggregation, and if it modulates ISR dysregulation in HD.

Modulation of the ISR has shown potential to influence HD phenotypes [22], with both ISR activation and inhibition found to be protective in HD. ISR activation counteracted disease phenotypes, namely reducing mut-Htt aggregation and promoting cell survival in cell models of N-terminal mut-Htt [23-25], and extending lifespan, increasing weight and improving motor performance in HD mice models expressing N-terminal mut-Htt (R6/2 and N171-82Q mice models) [26-28]. Conversely, ISR inhibition alleviated toxicity in HD cell models treated with ER stress-inducing agents [29, 30] and counteracted memory deficits and mut-Htt aggregation in the R6/1 HD mouse model [31]. In this study, we investigated if MPC inhibition could modulate the ISR activation and mitigate mut-Htt proteotoxicity in a HD model expressing N-terminal fragment of mut-Htt.

## Methods

### Cell Culture

PC12 cells with doxycycline-inducible expression of N-terminal fragment of the human wild-type (wt, 23 polyQ) or mutant (mut, 74 polyQ) huntingtin (Htt) fused to an enhanced green fluorescent protein (EGFP) at the N-terminus were obtained from Cambridge Enterprise – Lab of Dr. David C. Rubinsztein (Rub-1856-07, CL4 & CL5) [32]. Cell lines were cultured in standard conditions as we previously described [23]. Cells were seeded in multi-well plates coated with polyethyleneimine (PEI, Sigma-Aldrich, #P3143) and culture media was renewed every 48 h. Cells were seeded in 6-well plates at a density of 300,000 cells/cm^2^ and 120,000 cells/cm^2^ for ATP and protein extraction, respectively. For resazurin, medium pH and lactate dehydrogenase assays, cells were seeded in 96-well plates at a density of 135,000 cells/cm^2^. For mitochondrial respirometry assay, cells were seeded in XF24 cell culture microplates at a density of 300,000 cells/cm^2^. For live imaging assays, cells were seeded in 96-well imaging plates at a density of 60,000 cells/cm^2^.

### Drug treatments

The expression of the N-terminal fragment of wt- or mut-Htt was induced by 1 µg/mL doxycycline (Sigma-Aldrich, #D9891) 24 h after seeding, for 72 h [23]. To inhibit the MPC, cells were treated with UK5099 (Sigma-Aldrich, #PZ0160) and mitoglitazone (Sigma-Aldrich, #SML1884) simultaneously with doxycycline treatment. As positive controls for ISR activation, we used the ER stressor thapsigargin (Tocris, #1138, 0.01 µM [33]) and the activator of the ISR kinase PERK (protein kinase R-like ER kinase) CCT020312 (Millipore, #324879, 1 µM [23, 34]). As cellular stressors, we used the mitochondrial the complex I inhibitor rotenone (Sigma-Aldrich, #R8875, 0.5 µM [35] and the proteasome inhibitor MG-132 (SelleckChem, #S2619, 0.25 µM [36]).

Treatment with CCT020312 started in simultaneous with doxycycline treatment (induction). Treatment with thapsigargin, rotenone or MG-132 started 48 h after doxycycline treatment and was maintained for 24 h. Cell medium with drugs was renewed 48 h after induction. To measure culture medium pH, non-induced cells were treated with MPC inhibitors (0.1-100 µM) or rotenone, 24 h after seeding, for 72 h.

To measure mitochondrial respiration, non-induced cells were treated with MPC inhibitors immediately before starting the respirometry assay. In some of respirometry experiments, as specified in the corresponding figure legend, cells were supplemented with methyl pyruvate (Sigma-Aldrich, #371173, 10 mM) or assay medium after the basal readings. In each experiment, all conditions including the control condition presented the same percentage of drug solvent – maximum 0.6% DMSO (Sigma-Aldrich, #276855).

### Resazurin assay

Cell treatments were performed 24 h after seeding as described in Drug treatments section. Resazurin assay was performed as we previously described [23, 37]. After 68 h of drug treatments, 40 *μ*M resazurin (Sigma-Aldrich, #R7017) was added to each well and its reduction to resorufin was assessed by fluorescence readings every hour for 4 hours (completing 72 h of drug treatment). Results were expressed in percentage of solvent condition. Each condition was done in duplicate in each experiment.

### Medium pH values

Medium pH values were measured through ratiometric absorbance analysis of the pH indicator phenol red, as described previously [38]. Cells were seeded in the culture medium that contains phenol red (high glucose DMEM containing phenol red, Gibco, #11965092) and treated as described in Drug treatments section. After 72 h of treatment, 100 *μ*L of media from each well were transferred to a new 96-well plate and the absorbance of phenol red was measured immediately at 443 and 570 nm. Background signal was evaluated using culture medium with treatments in absence of cells. Results were expressed in absorbance 443 / 570 nm ratios – acidification of the medium increases the ratio. Each condition was done in triplicate in each experiment.

### Mitochondrial respiration

Mitochondrial oxygen consumption rate (OCR) was performed using a Seahorse XF24 extracellular flux analyzer (Seahorse Biosciences), following the manufacturer’s instructions. At 24 h before the assay, cells were plated on XF24 cell culture microplate and Seahorse probes were hydrated with Seahorse calibrant solution (pH 7.4). On the day of the assay, cells were washed twice with phosphate-buffered saline (PBS) and calibrated in a 37°C non-CO_2_ incubator with 500 μL of experimental medium (XF DMEM (Agilent, #103575-100) or DMEM (Sigma-Aldrich, #D5030) supplemented with 25 mM glucose (Sigma-Aldrich, #49159) and 4 mM GlutaMax (Gibco, #35050-038)) for 30 minutes. This medium was then replaced with fresh experimental medium containing the respective treatments as described in Drug treatments section. After measured basal OCR, 5 μM oligomycin B (ATP synthase inhibitor, Oligo, Sigma-Aldrich, #O-4876), 2 μM carbonyl cyanide-4-(trifluoromethoxy)phenylhydrazone (mitochondrial uncoupler to induce maximal respiration, FCCP, Sigma-Aldrich, #C-2920) and the combination of 1 μM rotenone and 1 µM antimycin A (inhibitors of complexes I and III, respectively, to induce a complete shutdown of the electron transport chain, Rot/Anti, Sigma-Aldrich, #A8674) were sequentially injected. In some of these experiments, 10 mM methyl pyruvate (MetPy, Sigma-Aldrich, #371173) was injected prior to the oligomycin B injection to rescue decreased OCR induced by MPC inhibitors [12]. Three OCR measurements were taken between each injection, following a protocol of 3 minutes mixing, 2 minutes waiting, and 3 minutes reading. At the end, the supernatant was removed, cells were lysed with sucrose lysis buffer [250 mM sucrose (Sigma-Aldrich, #84097), 20 mM HEPES (Sigma-Aldrich, #H4034), 3 mM EDTA (Sigma-Aldrich, #E6758); pH 7.5], and protein content quantified by standard Bradford protein assay (Bio-Rad, #500-0006). OCR values were normalized to protein content and to control baseline OCR. Each condition was done in duplicate in each experiment.

### ATP quantification

Intracellular ATP concentrations were quantified by high-performance liquid chromatography (HPLC) as previously described [39, 40] with some modifications. Cell treatments were performed as described in Drug treatments section. After 72 h treatment, culture medium was removed by aspiration, followed by immediate addition of 60 μL ice-cold 0.3 M perchloric acid per well. The plates were frozen at −80 °C for 10 minutes, followed by defrosting on ice for 15 minutes. Lysates were centrifuged at 3000 × g (10 min, 4 °C). Pellets containing acid-denatured proteins were stored at −80 °C for later protein quantification using Bradford protein assay, while supernatants were neutralized (pH 7) with 2 M KHO. Neutralized supernatants were allowed to rest for 30 min on ice and cleared by centrifugation at 14,000 × g (10 min, 4 °C) to deposit potassium perchlorate crystals. The resulting supernatants were filtered using 0.22 μm Nylon filter (Costar, #8169) immediately before the injection. ATP was quantified using HPLC (Thermo Scientific Dionex UltiMate 3000 Standard Systems) with a reversed-phase column [250 × 4.6 mm Luna 5 μm C18(2) 100 Å; Phenomenex], a diode-array detector (Dionex UltiMate 3000) and a computer with the Chromeleon CDS Software. The elution was performed in isocratic mode, with potassium phosphate buffer [60 mM K_2_HPO_4_ (Sigma-Aldrich, #1.05104), 40 mM KH_2_PO_4_ (Alfa Aesar, # A12142.36); pH 7] at 1.2 mL/min flux. Determinations were performed via 260 nm absorbance, using ATP (Sigma-Aldrich, #A7699) standard for calibration curves and specific peak identification. Results were expressed in μmol/g protein.

### Extracellular lactate dehydrogenase (LDH) activity assay

Extracellular LDH activity was measured as an indicator of cell death [41], based on the kinetics of NADH consumption kinetics linked to the conversion of pyruvate into lactate [23]. Cell treatments were performed as described in Drug treatments section. After 72 h of treatment, extracellular LDH and the total LDH (the extracellular and intracellular LDH) were quantified as we previously described [23]. Cytotoxicity was calculated as the ratio between extracellular LDH and total LDH content and presented as percentage of control.

### Protein extraction

Protein extraction from cells was performed as we previously described [23] with RIPA buffer and homogenization using Precellys Evolution after 3 cryofracture cycles. The homogenates were then centrifuged at 600 x g for 10 min at 4 °C. Protein was quantified in the supernatant by the Bradford protein assay.

### Western blotting

Western blot was performed as we previously described [23]. Denatured protein extracts (25 μg) were loaded into 4–12% Bis-Tris gels (Invitrogen, #NW04120BOX) or 4–15% TGX gels (BioRad, #456-1084) and electrophoresed at 200 V for 30 min. Then, proteins were wet-transferred to PVDF membranes (Merck, #IPVH00010) at 100 V for 60 min at 4 °C using a Mini Trans-Blot cell device (Bio-Rad). After blocking with 5% bovine serum albumin (BSA, NZYTech, #MB04602) in PBST [PBS with 0.05% Tween 20 (Sigma-Aldrich, #P9416)], membranes were incubated with primary antibodies, followed by the corresponding horseradish peroxidase-conjugated secondary antibodies. Detection was performed using a chemiluminescent kit (Novex ECL, Invitrogen, #WP20005) and imaged with a ChemiDoc MP Imaging system (Bio-Rad).

Membranes were re-probed with additional antibodies as needed. When the initial antibody targeted a protein with a similar molecular weight to the subsequent target, membranes were subjected to mild stripping. Briefly, membranes were washed with stripping buffer [200 mM glycine (NZYTech, #MB01401), 0.1% sodium dodecyl sulfate (SDS, Sigma-Aldrich, #L3771), 1% Tween 20; pH 2.2], PBS and PBST at room temperature with agitation, and re-blocked with 5% BSA in PBST prior to re-probing. This stripping protocol removes only the secondary antibody attached to the membrane; so, secondary antibodies used after stripping were selected from a different host species to avoid cross-reactivity. Stripping efficacy was confirmed using the chemiluminescent kit described above.

Coomassie staining of membranes was used for loading control. Densitometric analysis was performed with Image J (https://imagej.net/ij/). Briefly, background subtraction was applied to cropped blot regions containing the bands of interest. The integrated density of each band was then measured and normalized to the corresponding lane on the Coomassie-stained blot. Full blots are displayed in supplementary figures 1 to 4.

Primary antibodies: anti-CHOP - 1:1000, ProteinTech, #15204-1-A, RRID: AB_2292610; anti-eIF2α - 1:1000, Invitrogen, #AHO0802, RRID: AB_2536316; anti-GFP - 1:1000, Invitrogen, #MA5-15256, RRID: AB_10979281; anti-phospho-eIF2α - 1:500, Invitrogen, #MA5-15133, RRID: AB_10983400. Secondary antibodies: anti-mouse - 1:4000, Invitrogen, #G-21040, RRID: AB_2536527; anti-rabbit - 1:4000 anti-rabbit, Invitrogen, #G-21234, RRID: AB_2536530.

### Filter trap assay

Filter trap assay was performed as previously described [23], with some modifications. Briefly, protein extracts were diluted in 2% SDS in PBS to a final concentration of 0.3 μg/μL and denatured at 95 °C for 5 min. 100 μL of each sample (containing 30 μg of protein) was spotted in an acetate cellulose membrane (0.2 μm, Sterlitech, #CA022005) in duplicates, using a Bio-Dot Microfiltration Apparatus (Bio-Rad). To detect GFP-Htt aggregates, spotted membranes were blocked with 5% BSA in PBST and incubated with anti-GFP (1:1000, Invitrogen, #MA5-15256, RRID: AB_10979281) for 1 h, followed by incubation with horseradish peroxidase-conjugated anti-mouse (1:4000, Invitrogen, #G-21040, RRID: AB_2536527) for an additional hour. Detection was performed as described in the Western blotting section, and densitometric analysis of dots was performed using Image J. Full blots are displayed in supplementary figure 5.

### Live imaging

Cell treatments were performed as described in Drug treatments section. After 72 h of doxycycline treatment, live-cell imaging was performed using an inverted microscope (Eclipse TE300, Nikon), equipped with a motorized stage (ProScan, Prior), a monochromator (Polychrome II, Photonics), and a CCD camera (ORCA-ER, Hamamatsu), all controlled by the Micro-Manager 2.0 software [23, 42]. Hoechst 34580 (1 μg/mL, 1h, Sigma-Aldrich, 63493) was excited at 380 nm for nuclei visualization and EGFP-Htt at 488 nm for assessment of Htt expression. Emissions were collected using band pass filters (Chroma) for DAPI (Hoechst) and FITC (EGFP). Identical equipment settings were ensured between conditions and experiments.

### Statistical analysis

Data are plotted as mean ± standard error of the mean (SEM) of at least *n* independent experiments (*n* specified in figure legends). Concentration-response curves were fitted with non-linear regression – two phase decay or second order polynomial. The Shapiro-Wilk normality test was used to assess data normal distribution. For comparisons between two groups *t-test* was used for normally distributed data, while the Mann-Whitney’s test was used for non-normally distributed data. Single-factor analyses were performed using One-Way ANOVA followed by Dunnett’s or Sidak’s post-hoc tests. Two-Way ANOVA was used to test interactions and main effects between the treatments UK5099 and MG-132. For all analyses, a *P* value under 0.05 was taken as statistically significant. Data analyses were performed with Prism 9.0 (GraphPad).

## Results

### Mitoglitazone and UK5099 alter cellular metabolism by inhibiting the MPC

To evaluate the effects of MPC inhibition in a HD cellular model, we selected the pharmacological inhibitors UK5099 and mitoglitazone, as both have been described as effective and selective for the MPC [43-45]. They are representative of two different chemical classes, but both contain pyruvate-mimicking moieties that enable binding to the MPC [45]: UK5099 is an α-cyanocinnamate derivative with an IC_50_ of 50 nM for the MPC, which is 300-fold lower that the concentrations required for it to inhibit monocarboxylate transporters [44, 46], while mitoglitazone is a PPARγ-sparing thiazolidinedione: binding the MPC (IC_50_ = 1.3 µM) at 20-fold lower concentrations than those required for the PPARγ (23.7 μM) [43].

To characterize the concentration-dependent bioactivity of mitoglitazone and UK5099 in the HD cellular model we first measured their impact on resazurin metabolism. Cells expressing mut-Htt presented lower resazurin metabolism compared to cells expressing wt-Htt (Fig. 1a). Treatment with mitoglitazone and UK5099 up to 100 µM induced a concentration-dependent increase in resazurin metabolism in cells expressing either wt- or mut-Htt (Fig. 1a). However, at 300 µM, the highest tested concentration, both drugs decreased resazurin metabolism in comparison to control conditions (0 µM; solvent), suggesting cytotoxicity at 300 µM (Fig. 1a). Previous studies identified 10 µM UK5099 in primary neurons [47] and 10 µM mitoglitazone in differentiated neuronal precursor cells [4] as effective concentrations for MPC inhibition and induction of neuroprotective effects. Based on these previous studies and considering the observed decrease in resazurin reduction at 300 µM for both MPC inhibitors, we selected 10 and 100 µM concentrations of mitoglitazone and UK5099 for further analysis of MPC inhibition.

**Fig. 1.**
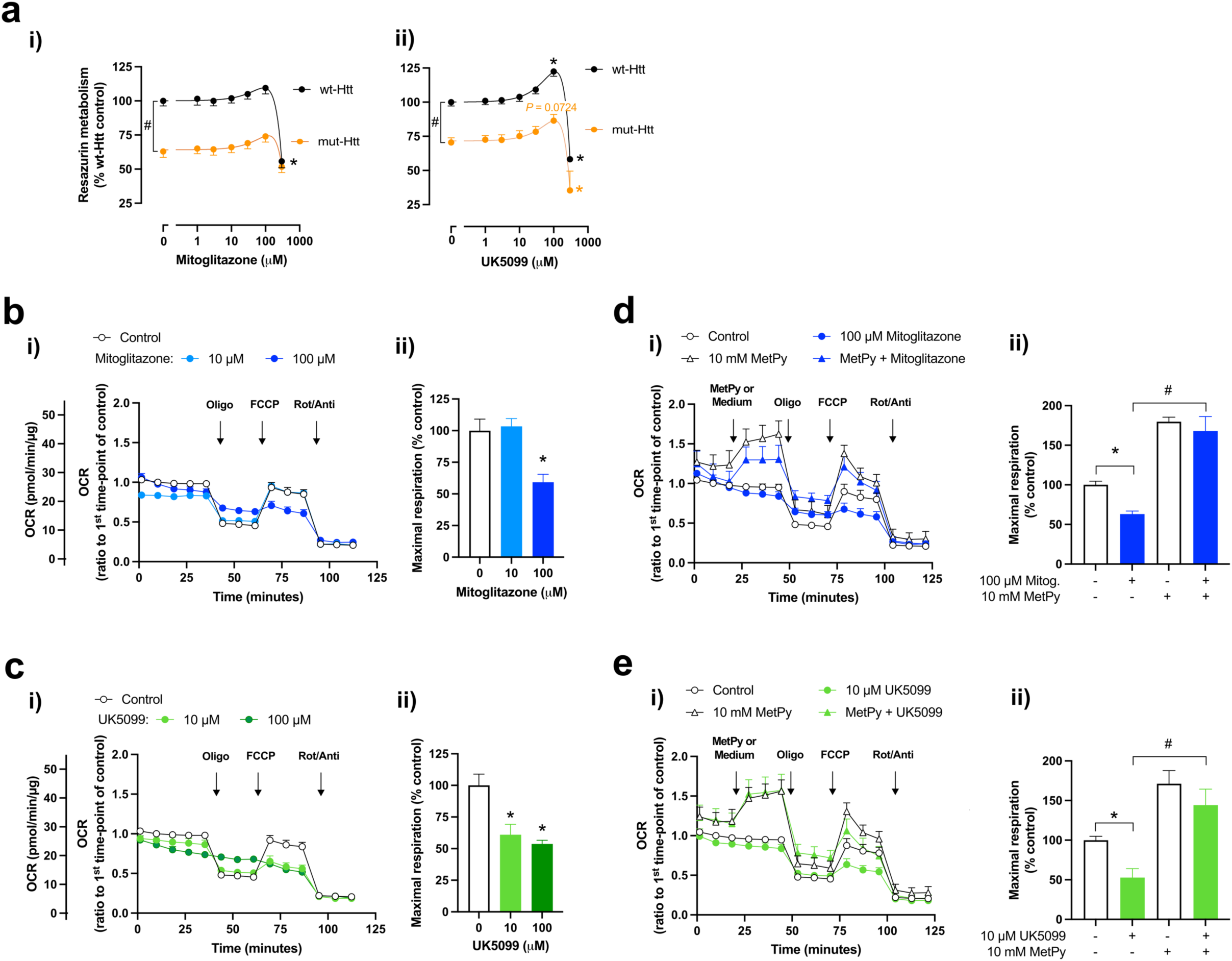
Effects of MPC inhibitors on cellular metabolism. **a** Resazurin metabolism of cells co-treated for 72 h with 1 µg/mL doxycycline – to induce the expression of wt-Htt or mut-Htt – and MPC inhibitors: (i) mitoglitazone (1-300 μM), *n* = 4 independent experiments; (ii) UK5099 (1-300 μM), *n* = 3-6 independent experiments. Data are mean ± SEM. ^#^*P* < 0.05, *t*-test; \**P* < 0.05 *vs*. respective control, One-Way ANOVA with Dunnett’s multiple comparison test. **b** Cells were treated with solvent (control) or mitoglitazone: (i) oxygen consumption rate (OCR) profile; (ii) maximal respiration; *n* = 2 - 4 independent experiments. **c** Cells were treated with solvent (control) or UK5099: (i) oxygen consumption rate (OCR) profile; (ii) maximal respiration; *n* = 3 - 4 independent experiments. **b, c** Data are mean ± SEM. \**P* < 0.05 *vs*. respective control, One-Way ANOVA with Dunnett’s multiple comparison test. **d** Cells treated with solvent (control) or mitoglitazone were challenged with 10 mM methyl pyruvate (MetPy), before addition of oligomycin (Oligo). Experimental medium was used as control of MetPy. (i) oxygen consumption rate (OCR) profile; (ii) maximal respiration; *n* = 2 independent experiments. **e** Cells treated with solvent (control) or UK5099 were challenged with 10 mM methyl pyruvate (MetPy), before addition of oligomycin (Oligo). Experimental medium was used as control of MetPy. (i) oxygen consumption rate (OCR) profile; (ii) maximal respiration; *n* = 2 independent experiments. **d, e** Data are mean ± SEM. \**P* < 0.05 *vs*. control, ^#^*P* < 0.05 vs. MPC inhibitor (mitoglitazone or UK5099), One-Way ANOVA with Sidak’s multiple comparison test. Oligo, oligomycin B (5 μM); FCCP, carbonyl cyanide-4-(trifluoromethoxy)phenylhydrazone (2 μM); Rot, rotenone (1 μM); Anti, antimycin A (1 μM)

To verify if the selected concentrations of mitoglitazone and UK5099 inhibit the MPC, we tested whether they reduced the mitochondrial oxygen consumption rate (OCR), and if such reduction could be rescued by a membrane permeable pyruvate analogue. The rationale is that MPC inhibition, by limiting the mitochondrial pyruvate uptake, should reduce substrate availability for the TCA cycle and the respiratory chain, consequently decreasing oxygen consumption [12, 48]. Mitoglitazone at 100 µM reduced OCR (Fig. 1bi), as evidenced by the reduced maximal respiration (Fig. 1bii). UK5099 at both 10 and 100 µM also decreased maximal respiration (Fig. 1c). Administration of the membrane-permeable methyl pyruvate [12] increased basal and maximal respiration, rescuing the OCR reduction caused by mitoglitazone (Fig. 1d) and UK5099 (Fig. 1e). These results indicate that the effects of mitoglitazone and UK5099 on the OCR, particularly on maximal respiration, are due to MPC inhibition and the consequent limitation of pyruvate entry into mitochondria.

### MPC inhibitors attenuate increased ISR activation in the HD cell model

To investigate the hypothesis that MPC inhibition modulates the ISR in a HD cellular model, we quantified the levels of key ISR effectors after MPC pharmacological inhibition.

In the absence of drug treatments, cells expressing mut-Htt presented increased levels of phosphorylated eIF2α (p-eIF2α), but reduced levels of CHOP compared to cells expressing wt-Htt (Fig. 2; # *p*<0.05), suggesting that mut-Htt expression induces an ISR activation with suppressed CHOP induction.

**Fig. 2.**
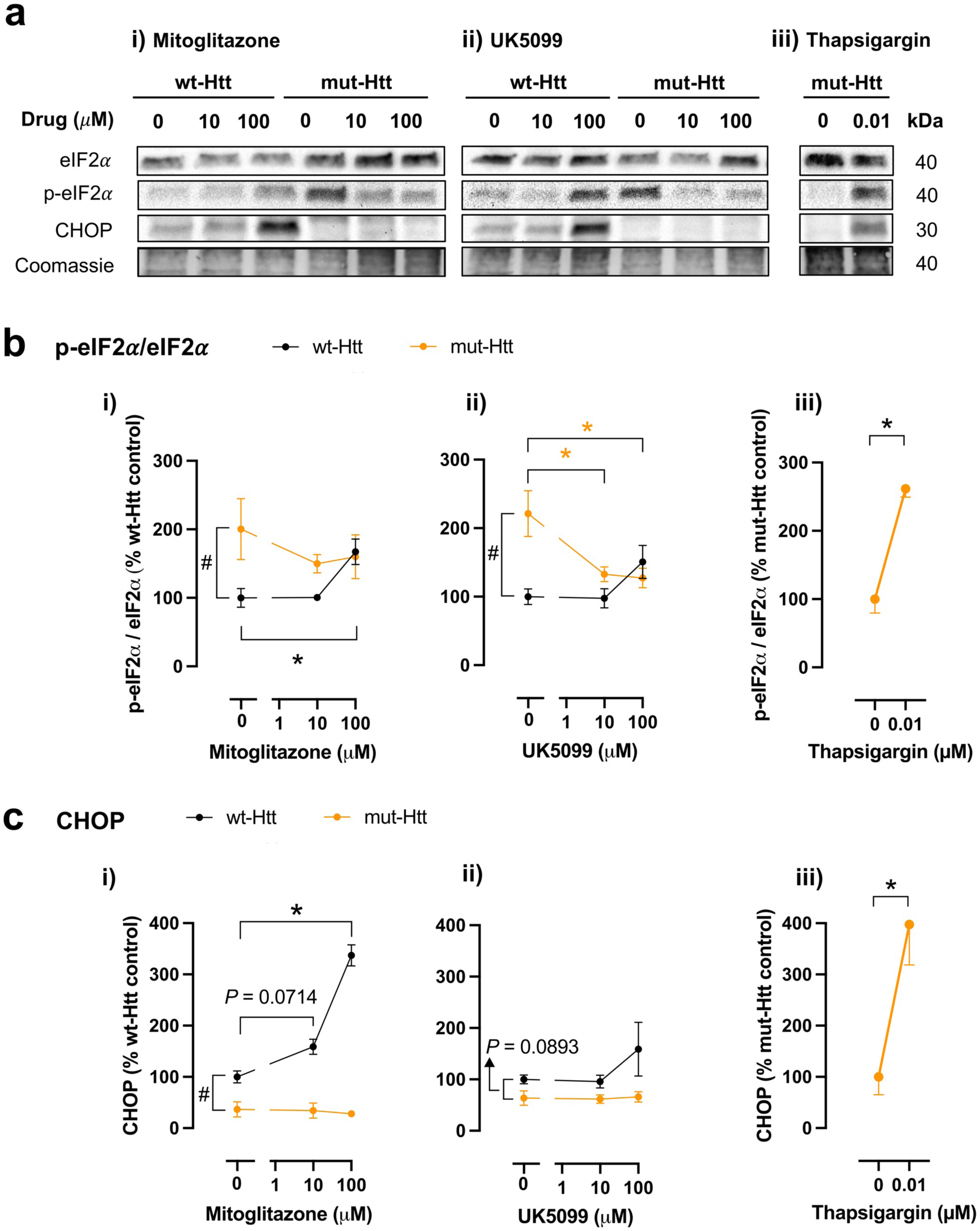
Effect of MPC inhibitors on the ISR pathway. Cells were co-treated for 72 h with doxycycline (1 µg/mL) – to induce the expression of wt-Htt or mut-Htt – and solvent (control) or MPC inhibitors: mitoglitazone and UK5099. Thapsigargin (0.01 μM), an ISR inducer, was used as positive control of the ISR activation. **a-c** Representative western blots of the ISR markers (**a**) and respective quantification of p-eIF2α/eIF2α (**b**) and CHOP levels (**c**) in cells expressing wt- or mut-Htt, treated with: mitoglitazone (i); UK5099 (ii); or thapsigargin (iii). Data are mean ± SEM from 3-4 independent experiments. ^#^*P* < 0.05, *t*-test comparison between wt-Htt- and mut-Htt-expressing cells; \**P* < 0.05 *vs*. respective control: One-Way ANOVA with Dunnett’s multiple comparison test for mitoglitazone (i) and UK5099 (ii) data; *t*-test for thapsigargin data (iii)

In cells expressing wt-Htt, treatment with 100 µM mitoglitazone significantly increased p-eIF2α (Fig. 2bi) and CHOP levels (Fig. 2ci) compared to solvent-treated cells. Similarly, treatment with 100 µM UK5099 showed a trend for increased p-eIF2α (Fig. 2bii) and CHOP levels (Fig. 2cii).

In cells expressing mut-Htt, treatment with 10 and 100 µM UK5099 significantly decreased p-eIF2α compared to solvent-treated cells (Fig. 2bii). Similarly, 10 and 100 µM mitoglitazone showed a trend toward decreased p-eIF2α (Fig. 2bi). Regarding CHOP levels, neither mitoglitazone nor UK5099 altered CHOP levels in cells expressing mut-Htt (Fig. 2ci-ii).

Given the low basal CHOP levels found in cells expressing mut-Htt (Fig. 2ci-ii), we tested whether these cells retained the capacity to further activate the ISR and upregulate CHOP expression upon treatment with the ER stressor thapsigargin [33]. Thapsigargin at 10 nM significantly increased both p-eIF2α and CHOP levels in cells expressing mut-Htt (Fig. 2a-ciii). These results show that these cells retain the capacity to induce CHOP expression in response to a potent stimulus like thapsigargin, which disrupts calcium homeostasis and activates ISR by inducing ER stress [49].

Altogether, these results demonstrate differential effects of MPC inhibitors on ISR activation depending on the presence of mut-Htt: while MPC inhibition activates the ISR pathway in cells expressing wt-Htt, it attenuates the ISR activation induced by mut-Htt.

### MPC inhibitors do not compromise cellular ATP levels

ISR activation optimizes the use of available energy resources [49]. Such energetic optimization should be essential when the MPC is inhibited and thereby limits the supply of pyruvate to mitochondria [6, 7]. Given that MPC inhibitors attenuated ISR activation in mut-Htt-expressing cells (Fig. 2bi, ii), we hypothesised that MPC inhibitors might negatively impact the cellular energetic status. We tested this hypothesis by comparing the ATP levels of cells with vs. without treatment with MPC inhibitors, using rotenone as a positive control for energetic impairment [50].

Cells expressing mut-Htt presented lower ATP levels than those expressing wt-Htt (Fig. 3a), consistent with their reduced resazurin metabolism (Fig. 1a). Treatment with the positive control rotenone reduced cellular ATP levels (Fig. 3ai). Treatment with mitoglitazone significantly increased ATP levels in mut-Htt-expressing cells and showed a trend for increased ATP levels in wt-Htt-expressing cells (Fig. 3aii). Treatment with UK5099 significantly increased ATP levels only in wt-Htt-expressing cells maintaining the ATP levels in mut-Htt-expressing cells (Fig. 3aiii). These results indicate that these concentrations of MPC inhibitors do not compromise the cellular energetic status even in the presence of mut-Htt.

**Fig. 3.**
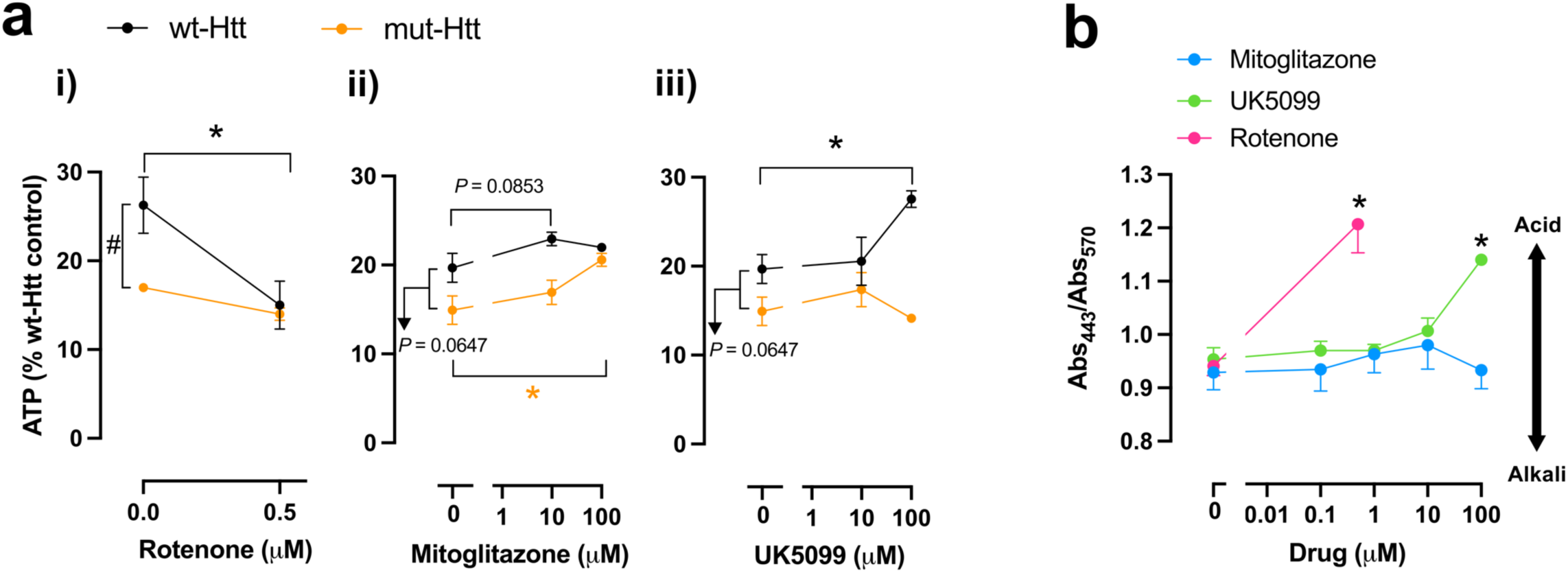
Effect of MPC inhibitors on ATP levels and extracellular acidification. **a** ATP levels of cells co-treated for 72 h with doxycycline (1 µg/mL; to induce the expression of wt-Htt or mut-Htt) and the mitochondrial complex I inhibitor rotenone (0.5 μM; positive control for effects on ATP) (i) or MPC inhibitors: mitoglitazone (ii) or UK5099 (iii). Data are mean ± SEM from 3 independent experiments. ^#^*P* < 0.05, *t*-test comparison between wt-Htt- and mut-Htt-expressing cells; \**P* < 0.05 *vs*. respective control: *t*-test for rotenone data (i); One-Way ANOVA with Dunnett’s multiple comparison test for mitoglitazone (ii) and UK5099 (iii). **b** pH variation of the culture media of cells treated with solvent (control) or MPC inhibitors (mitoglitazone and UK5099). Rotenone (0.5 μM) was used as positive control for extracellular acidification. pH was measured based on the ratiometric property of the pH indicator, phenol red, and presented as a ratio between the absorbances at 443 nm and 570 nm. Data are mean ± SEM from 3-6 independent experiments. \**P* < 0.05 *vs*. respective control: *t*-test for rotenone data (i); One-Way ANOVA with Dunnett’s multiple comparison test for mitoglitazone (ii) and UK5099 (iii)

To further complement the study of metabolic alterations upon MPC inhibition, we studied, in non-induced cells, the effects of MPC inhibition on extracellular acidification, which can be associated with a higher conversion of pyruvate to lactate by lactate dehydrogenase [51]. We used rotenone (0.5 µM) as a positive control for medium acidification (Fig. 3b), since it increases glycolysis and lactate secretion by inhibiting mitochondrial complex I [35]. Mitoglitazone up to 100 µM maintained the culture medium acidification similar to the control condition (Fig. 3b), while UK5099 at 100 µM increased acidification of the culture medium (Fig. 3b). While we did not find an increase of extracellular acidification induced by mitoglitazone treatment, our data show that UK5099 has a significant impact on extracellular acidification. This effect may result from pyruvate that cannot enter the mitochondria due to MPC inhibition and is subsequently converted to lactate.

### MPC inhibitors do not reduce N-terminal mut-Htt levels or cell death

To investigate if the MPC inhibitors could alter mut-Htt proteostasis, we monitored the aggregation phenotype (Fig. 4a, b), and quantified SDS-insoluble Htt aggregates and soluble Htt levels, respectively, by filter trap and Western blotting assays. As positive controls for modulating mut-Htt aggregation, we used the proteasome inhibitor MG-132 (0.25 µM) [52] and the ISR kinase PERK activator CCT020312 (1 µM) [23, 34]. MG-132 significantly increased the levels of mut-Htt aggregates (Fig. 4ciii, d), while CCT020312 significantly decreased them (Fig. 4ciii, d), demonstrating that mut-Htt proteostasis is pharmacologically modulable in this HD cell model.

**Fig. 4.**
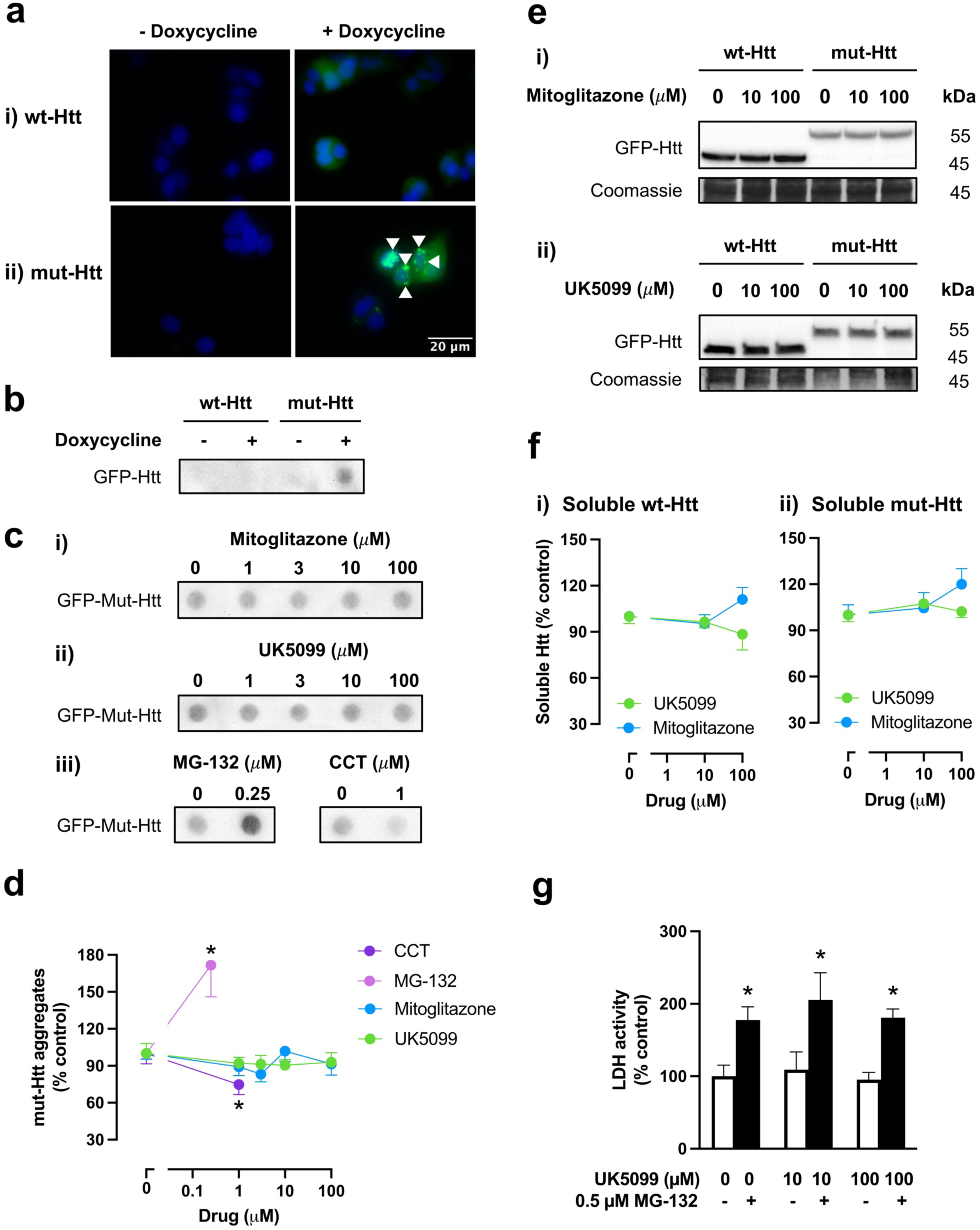
MPC inhibitors and Htt proteostasis. Cells were co-treated for 72 h with doxycycline (1 µg/mL) to induce the expression of wt-Htt or mut-Htt, and solvent (control) or MPC inhibitors: mitoglitazone and UK5099. MG-132 (0.25 μM; a proteasome inhibitor) and CCT020312 (1 μM; an ISR kinase activator) were used as positive controls for the modulation of mut-Htt aggregation. **a** Representative images of cells expressing wt- (i) or mut-Htt (ii); Htt expression (EGFP, green), nucleus staining (Hoechst 34580, blue). Cells expressing mut-Htt (+ doxycycline) form aggregates (white arrowheads). **b** Representative blot from filter trap assay of Htt aggregation in cells expressing wt- or mut-Htt without and with doxycycline treatment. **c** Representative blots (from filter trap assay) of mut-Htt aggregates in cells expressing mut-Htt treated with: mitoglitazone (i); UK5099 (ii); MG-132 or CCT020312 (iii). **d** Quantification of mut-Htt aggregates shown in **c**i-iii. Data are mean ± SEM from 3-8 independent experiments. Mann-Whitney test (MG-132) or *t*-test (CCT020312): **\****P* < 0.05 *vs.* respective control. **e** Representative western blots of soluble wt- and mut-Htt of cells treated with: mitoglitazone (i) or UK5099 (ii). **f** Quantification of soluble wt- (i) and mut-Htt (ii) shown in (**e**). Data are mean ± SEM from 4 independent experiments. **g** Extracellular lactate dehydrogenase (LDH) activity as a measure of cell death of mut-Htt expressing cells treated with UK5099 in the absence or in presence of 0.25 µM MG-132. Data are mean ± SEM from 3-6 independent experiments. Two-Way ANOVA, \**P* < 0.05, MG-132 effect

Treatment with the MPC inhibitors mitoglitazone or UK5099 had no significant impact in either mut-Htt aggregation (Fig. 4c, d), or in the levels of soluble Htt compared to solvent control (Fig. 4e, f). Still, given the robust effect of UK5099 in attenuating ISR activation in mut-Htt-expressing cells (Fig. 2bii), we tested whether UK5099 could reduce cell death in this cell model. To evoke significant cell death, we used the co-stressor MG-132, which is a proteasome inhibitor, reducing mHtt-clearance and enhancing cell death. We show that MG-132 (0.25 µM) enhanced N-terminal mut-Htt aggregation (Fig. 4ciii, d) and increased cell death assessed by the LDH release assay (Fig. 4g). However, co-treatment with UK5099 (10-100 µM) did not rescue the increased cell death (Fig. 4g).

Taken together, these results show that mut-Htt proteostasis in this cell model can be pharmacologically modulated (e.g. by proteasome inhibition or PERK activation), but the MPC inhibitors – despite attenuating ISR activation – were unable to reduce aggregation or rescue cell death evoked by high levels of N-terminal mut-Htt.

## Discussion

MPC inhibition was neuroprotective in models of neurodegenerative disorders such as PD and AD [2-5]. However, the neuroprotective role of MPC inhibition in HD, particularly regarding mut-Htt proteostasis, remains mostly unexplored. Indeed, current knowledge of the effects of MPC inhibition on proteostasis – typically compromised in neurodegenerative disorders is limited to findings in AD models, where MPC knockout or inhibition reduced Aß and tau aggregates [2]. Here, we investigate the effects of MPC inhibition in a cellular model of HD expressing the N-terminal fragment of mut-Htt, which presents proteotoxic stress, metabolic impairment and overactivation of the ISR [22, 23], a pro-survival pathway that regulates both metabolism and proteostasis [9].

The ISR integrates different cellular stress signals, such as mitochondrial dysfunction, amino acid deprivation and proteotoxicity [53]. Here, we show that MPC inhibition activates the ISR under control conditions (in wt-Htt-expressing cells). Similar ISR activation after MPC inhibition was previously found in human hair follicles and in cellular models of prostate and lung cancer [11-13]. The metabolic reprogramming induced by MPC inhibition may explain the ISR activation: MPC inhibition reduces mitochondrial pyruvate import and increases amino acid catabolism [6, 7]; the resulting reduction of cytosolic amino acid pool activates the ISR kinase GCN2, stimulating the ISR pathway [10]. The ISR activation is essential for preserving energy levels and proteostasis by reducing global protein synthesis and activating autophagy [9, 53].

Mut-Htt activates the ISR, as shown in this work and supported by previous findings in HD models and patient samples [23, 24, 29, 54-56]. The HD cell model with N-terminal mut-Htt used in this work presented ISR activation with increased p-eIF2α and reduced CHOP levels. A previous study has also reported reduced CHOP levels in immortalized striatal-like cells expressing full-length mut-Htt HdhQ111 [57], potentially associated with decreased levels of the mitochondrial transporter ABCB10: ABCB10 levels were found reduced in the striatum of HD R6/2 mice and in fibroblasts of HD patients and its deletion significantly decreased CHOP levels in the control cell line HdhQ7 cells (immortalized striatal-like cells expressing full-length wt-Htt) [57, 58].

Here we provide original evidence that MPC inhibitors can attenuate mut-Htt-induced ISR activation. These findings are consistent with related studies showing that MPC deletion attenuated ISR activation in cells exposed to the ER stressors tunicamycin and thapsigargin, reducing CHOP levels [14]. But what mechanisms could explain how MPC inhibition attenuates ISR activation induced by a cellular stressor such as mut-Htt? In HD, mitochondrial dysfunction (evidenced here by reduced resazurin metabolism and ATP levels in mut-Htt-expressing cells compared to wt-Htt-expressing cells) may impair NADH consumption by oxidative phosphorylation [59], leading to elevated NADH/NAD^+^ ratios, as observed in the striatum of HD mice [60]. Elevated NADH/NAD^+^ may activate the ISR [61], by hindering aspartate synthesis and depleting asparagine, which activates the ISR kinase GCN2 [62]. Thus, since MPC inhibition diverts pyruvate to lactate production, consuming NADH instead of producing it [63], we propose that MPC inhibition reduces the elevated NADH/NAD^+^ that is activating the ISR in mut-Htt-expressing cells and, consequently, MPC inhibition attenuates their ISR activation.

ISR activation is commonly observed in neurodegeneration, and ISR inhibition has been proposed as a potential therapeutic strategy for several neurodegenerative disorders [22, 64]. Thus, the attenuation of the ISR by MPC inhibition could explain the protective effects of MPC inhibitors observed in previous studies in AD and PD. Additionally, previous studies suggest that MPC inhibition decreases acetyl-coenzyme A levels, inhibiting the nutrient-sensing mammalian target of rapamycin mTOR pathway and increasing autophagy [3, 4, 65]. Autophagy plays a crucial role in maintaining cellular proteostasis by removing misfolded and aggregated proteins or dysfunctional organelles [66], further supporting a role of MPC inhibition in proteostasis regulation.

As HD is a monogenic neurodegenerative disorder caused by the expression of mut-Htt, a protein prone to undergo proteolysis, misfolding and aggregation [18], it is an excellent model disease to test experimental therapeutic strategies on proteostasis. Here, we studied the effects of MPC inhibition on the proteostasis of a cellular HD model expressing N-terminal mut-Htt. We also examined the impact of MPC inhibition on cell viability of this HD model, which required co-stressing cells with the proteasome inhibitor MG-132 to potentiate mut-Htt aggregation and evoke significant cell death. We showed that MPC inhibition did not change levels or aggregation of mut-Htt, nor did it reduce cell death in mut-Htt-expressing cells co-stressed with MG-132. Thus, although ISR modulation might contribute to the neuroprotective effects of MPC inhibition, our findings suggest that MPC inhibition alone may not be sufficient to improve proteostasis or prevent cell death in a model characterized by rapid N-terminal mut-Htt aggregation.

In conclusion, aiming to investigate the role of MPC inhibitor on ISR modulation and its potential protective effects in a HD cell model, we evaluated the ISR activation, metabolic adaptations and mut-Htt proteostasis following MPC inhibition. We show that MPC inhibitors differentially modulate the ISR activation depending on the presence of mut-Htt: MPC inhibitor activates the ISR under control conditions (in the presence of wt-Htt), while attenuating ISR activation induced by mut-Htt. However, our results also show that MPC inhibitors may not effectively reduce mut-Htt levels and aggregation or prevent cell death in models expressing N-terminal mut-Htt, which are characterized by a rapid and severe disease progression. Despite this, MPC inhibitor may be a potential therapeutic strategy for fine-tuning the ISR and warrant further investigation in models expressing full-length mut-Htt, which exhibit slower disease progression.

## Abbreviations

AD: Alzheimer’s disease
ATF4: activating transcription factor 4
CHOP: C/EBP homologous protein
eIF2*α*: eukaryotic translation initiation factor 2*α*
ER: endoplasmic reticulum
GCN2: non-depressible general control 2
HD: Huntington’s disease
Htt: huntingtin
ISR: integrated stress response
LDH: lactate dehydrogenase
MetPy: methyl pyruvate
MPC: mitochondrial pyruvate carrier
Mut-Htt: mutant Htt
p-eIF2*α*: phosphorylated eIF2*α*
OCR: oxygen consumption rate
PD: Parkinson’s disease
PPARγ: peroxisome proliferator-activated receptor gamma
TCA: tricarboxylic acid
Wt: wild-type.

## Statements and Declarations

### Author contributions

Ângela Oliveira: Conceptualization, Methodology, Validation, Formal analysis, Investigation, Data Curation, Writing - Original Draft, Writing - Review & Editing, Visualization. Liliana M. Almeida: Methodology, Validation, Formal analysis, Investigation. Brígida R. Pinho: Conceptualization, Methodology; Formal analysis, Data Curation, Writing - Original Draft; Writing - Review & Editing, Supervision. Jorge M. A. Oliveira: Conceptualization, Methodology; Resources, Writing - Review & Editing, Supervision, Project administration, Funding acquisition.

### Data Availability

The data that support the findings of this study are available from the corresponding authors upon reasonable request.

